# CryoConvNeXt enables robust identification of low-abundance molecular species in experimental cryo-EM data

**DOI:** 10.64898/2026.09.08.750038

**Authors:** Larissa Glass, Jan Pieter Abrahams, Thomas Braun

**Affiliations:** Biozentrum, University of Basel, 4056 Basel, Switzerland; Laboratory of Nanoscale Biology, Paul Scherrer Institut 5232 Villigen PSI, Switzerland; Laboratory of Biological Electron Microscopy, Institute of Physics, School of Basic Science, École Polytechnique Fédérale de Lausanne (EPFL), 1015 Lausanne, Switzerland

## Abstract

Rare but biologically important molecular species are easily lost when reconstruction-based cryo-EM classification is applied to heterogeneous samples dominated by more abundant particles. We developed CryoConvNeXt, a deep-learning classifier that combines cyclic equivariance with adaptive frequency filtering. It is trained on simulated projections and adapted to experimental data by self-training, an unsupervised domain adaptation technique where the model acts as its own annotator. We tested CryoConvNeXt on cryo-EM data collected for this study from controlled binary and ternary mixtures of Catalase, Apoferritin, and HSP60. Manual particle curation provided reference labels for benchmarks with class ratios from 1:1 to 1:16. CryoConvNeXt retained minority-species recall across this range. In contrast, cryoSPARC’s recall collapsed in four of six pairwise conditions, as early as 1:4 when Catalase was the minority species. By recovering low-abundance species that conventional classification fails to recover reliably, CryoConvNeXt takes an important step towards solving a central problem in quantitative visual proteomics: measuring the molecular composition of complex biological samples.

## 1 Introduction

Electron microscopy of vitrified specimens (cryo-EM) has experienced a renaissance as a tool in structural biology. Its single-particle sensitivity and minimal sample requirements make it, in principle, an ideal analytical method for studying heterogeneous mixtures directly from nanoliter-scale samples. In practice, however, the inherently low signal-to-noise ratio of particle projections has long limited reliable classification of heterogeneous species. Recent advances in deep-learning-based image analysis are beginning to change this (Levy *et al*., 2024*a*; Punjani & Fleet, 2021; Chen & Ludtke, 2021), opening new routes to quantitative analysis of compositionally complex biological samples.

Despite these advances, the analysis of heterogeneous samples remains a central challenge in single particle analysis (SPA) (Kimanius & Schwab, 2024). Biological preparations routinely contain multiple coexisting species – multimers, assembly intermediates, co-purified contaminants, and damaged particles among them – making classification a prerequisite for quantitative compositional analysis.

Structural heterogeneity is a fundamental property of biological systems: macromolecular assemblies coexist and interconvert in non-equilibrium steady states, and their relative abundances encode functional states. Characterising this structural proteome, i.e. the assemblies present in a cell and their relative populations, is therefore central to understanding molecular mechanism, and disease in particular. In neurodegenerative disorders, for instance, it is the distribution of coexisting assembly states of amyloidogenic proteins that determines pathological outcome, rather than their identity alone. Cryo-EM has the potential to read out these states directly through visual proteomics, in which macromolecular complexes are identified and structurally characterised within heterogeneous cellular extracts by SPA-like approaches (Arnold *et al*., 2018).

However, this goal introduces an additional challenge: the abundance distribution of macromolecular species in endogenous samples is often characterised by severe species imbalance. While abundant complexes may contribute hundreds of thousands of particle projections, rare but biologically important species may appear only in a small fraction of the data, with differences in relative abundances spanning several orders of magnitude (Morgenstern *et al*., 2021).

Standard cryo-EM classification algorithms, such as those based on Expectation-Maximisation heterogeneous reconstruction (Grant *et al*., 2018; Scheres *et al*., 2005; Scheres, 2012), often exhibit a bias toward high-abundance particles. In these scenarios, rare particles are frequently misclassified as noise or absorbed into the conformational averages of the majority species, effectively rendering the minority species invisible. To resolve these rare species, current workflows require the collection of massive datasets and often extensive semi-manual pruning to ensure the minority signal exceeds the threshold of detection.

In earlier work, we introduced differential visual proteomics (Syntychaki *et al*., 2019), a purely analytical framework that applies principal component analysis (PCA) to 2D class averages to detect structural differences between biological conditions in an unbiased manner. This revealed, for instance, the upregulation of heat-shock proteins without any targeted intervention. However, as the underlying 2D classification relies on Expectation-Maximisation, it inherits the majority-species bias discussed above, limiting sensitivity in the case of rare or low-abundance components.

A separate line of work targets heterogeneity within a single dataset through 3D reconstruction: neural ab initio reconstruction (Levy *et al*., 2024*a*), 3D variability analysis (Punjani & Fleet, 2021), and Gaussian-mixture models coupled to a neural network (Chen & Ludtke, 2021). These methods are optimised for reconstruction quality across a continuous heterogeneity landscape and discrete classes emerge only from post hoc clustering. Thus, rare or low-abundance states can be easily missed.

Here, we address the problem of multi-particle differentiation for cryo-EM datasets with pronounced species imbalance by employing a neural network classifier, CryoConvNeXt. The optimal training set for any supervised classifier would be labelled experimental data drawn from the same distribution as the images to be classified. Yet this is impractical, as obtaining reference particle labels through e.g. semi-manual curation of heterogeneous reconstruction outputs is laborious. Furthermore, rare species may contribute too few particles to train on reliably. We therefore train CryoConvNeXt on simulated data, for which unlimited labelled projections can be generated automatically for any species with a known density map. Training on simulated rather than experimental images, however, introduces a simulation-to-real domain gap. We narrow this gap by calibrating the simulation to the specific microscope and detector used for data collection, including MTF, CTF, and noise characteristics. The remaining gap is further narrowed through self-training, an unsupervised domain adaptation technique that incorporates high-confidence experimental predictions into the training set over successive rounds, adapting the model to the experimental imaging characteristics before inference.

We designed and collected an experimental cryo-EM benchmark dataset of three structurally distinct proteins and manually generated reference labels. We evaluate CryoConvNeXt on this dataset and benchmark its performance against cryoSPARC heterogeneous reconstruction. CryoSPARC was selected because it represents the most widely used heterogeneous reconstruction workflow and provides a strong practical baseline for routine cryo-EM processing. CryoConvNeXt trained on simulated data matches cryoSPARC under balanced conditions. With self-training it outperforms cryoSPARC, especially under species imbalance. CryoConvNeXt recovers minority species even where cryoSPARC’s heterogeneous reconstruction fails to resolve them.

More broadly, these results establish simulation-to-real transfer learning as a practical route to classification in cryo-EM, and position CryoConvNeXt as one building block of quantitative visual proteomics.

## 2 Methods

### 2.1 Neural Network Architecture and Training

We introduce CryoConvNeXt: a neural network architecture that consists of four sequential components: a learnable Butterworth filter, a C4-equivariant convolutional stem, ordinary ConvNeXt blocks and a radial attention gate. The Butterworth filter adaptively suppresses high spatial frequencies, reducing the contribution of noise-dominated frequency bands. The equivariant stem applies a shared 7 *×* 7 kernel rotated explicitly to 0^°^, 90^°^, 180^°^, and 270^°^, and fuses the resulting feature maps via max-over-group pooling to produce representations that are invariant to in-plane rotations by multiples of 90^°^. This exploits the fact that in cryo-EM, the in-plane orientation of each particle is a nuisance variable. The subsequent ConvNeXt blocks, a modernised ResNet incorporating larger kernels and depthwise convolutions, can therefore learn discriminative features without needing to account for in-plane rotations by multiples of 90^°^. The radial attention gate, inserted between the second and third ConvNeXt stages, allows the network to up-weight informative frequency bands and attenuate others.

We use a label-smoothing cross-entropy loss function with *ϵ* = 0.1 to mitigate overfitting, thereby reducing overconfident predictions and improving generalisation to out-of-distribution data. This means that instead of targeting a confidence of 1 for the correct class and 0 for the incorrect classes, we target a confidence of 0.9 for the correct class and distribute the remaining 0.1 uniformly across the incorrect classes.

ConvNeXt blocks are off-the-shelf (Liu *et al*., 2022), as are label-smoothing (Szegedy *et al*., 2016), cross-entropy loss and the AdamW optimizer (Loshchilov & Hutter, 2019). The cyclic equivariant stem is a simplified version of the group equivariant stem (Cohen & Welling, 2016). The learnable Butterworth filter is implemented as a custom layer in PyTorch, adapted from the Butterworth filter implemented in the scikit-image library (van der Walt *et al*., 2014), with the cutoff frequencies and the order of the filter as learnable parameters. The radial attention gate is a custom layer. It bins each feature map’s 2D power spectrum into radial frequency shells, passes that per-image radial profile through a small MLP to produce a per-channel sigmoid gate, and rescales the feature map channel-wise, letting the network attenuate frequency bands corrupted by CTF zeros or noise.

### 2.2 Simulated Data

To provide an unlimited supply of labelled training data for CryoConvNeXt, we use simulated data. This requires 3D density maps as input but provides in return a fully automated, unlimited, and labelled training set. We use the TEM-simulator (Rullgård *et al*., 2011) to generate a library of simulated 2D projections from 3D density maps for each species in our mixtures, sampling uniformly over the full range of particle orientations.

The TEM-simulator models the ideal exit wave intensity using Phase Object Approximation and subjects it to primary noise by Poisson sampling to simulate the stochastic nature of electron detection. It is then subjected to secondary noise, which is modelled using an Inverse Gaussian (Wald) distribution to account for variability in signal amplification and noise characteristics of the detector. Finally, the noisy detector response is convolved with the detector’s point spread function to simulate detector blurring.

We simulate ~9000 micrographs across all mixture conditions; full simulation parameters are provided in Table 3 (Appendix). These micrographs are imported into cryoSPARC (Punjani *et al*., 2017), where CTF estimation is performed with ctffind4 (Rohou & Grigorieff, 2015) and exposures are curated at a CTF fit resolution cutoff of 15 Å. Particles are picked with Topaz (Bepler *et al*., 2019) using a pretrained model (ResNet16 32 units), an estimated particle diameter of 160 Å, and an extraction threshold (log likelihood score) of 1. This processing pipeline mirrors the experimental workflow as closely as possible, minimising the simulation-to-real domain gap. We generated a particle stack of 70 000 particles per species and ~40 000 empty picks.

### 2.3 Self-Learning to Improve Transfer Learning

To improve transfer learning from simulated to experimental data, we employ self-training, a method for unsupervised domain adaptation, i.e. adapting from the simulated to the experimental domain. After training the model on simulated data, we evaluate the experimental test set and identify the top 25% of predictions based on confidence scores. These high-confidence predictions are then incorporated into the simulated training set for the next round of training. This process is repeated, incorporating the top 50% of predictions into the simulated data, allowing the model to iteratively refine its understanding of the experimental data distribution and improve its performance. This process is fully automated: no manual intervention or additional labelling is required. The method is transductive, adapting to the target data distribution before inference.

### 2.4 Evaluation Metrics

Standard classification accuracy metrics weigh all samples equally. This can be deceptive for imbalanced datasets, as for instance at high imbalance a naive classifier that consistently predicts the majority class will yield a high score, while completely failing to identify the minority class. As we are interested in the performance on minority classes, we instead evaluate our model using the macro-averaged *F*_1_ score (*F*_1,macro_), which treats each class equally regardless of its frequency in the dataset. The *F*_1,macro_ score is calculated as follows:

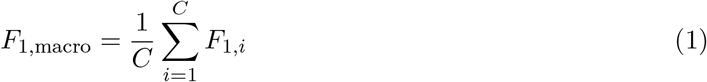

where *F*_1,*i*_ is the individual *F*_1_ score for class *i*, and *C* is the total number of classes. The class-specific *F*_1,*i*_ is defined as:

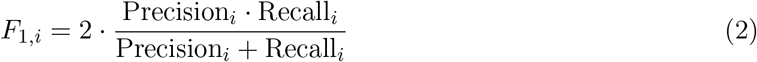

Precision represents the fraction of true positive assignments relative to all images classified as a given class (protein), quantifying the cleanliness of the isolated population. Recall, conversely, denotes the fraction of true positives retrieved from the total pool of actual instances of that protein, measuring the completeness of the extraction.

To complement the macro-averaged framework in the case of imbalanced scenarios, we explicitly track the individual *F*_1_ score, precision and recall for the minority species.

Furthermore, we evaluate Receiver Operating Characteristic (ROC) curves and the corresponding Area Under the Curve (AUC) for each class in the balanced (1:1) Apoferritin–Catalase, HSP60– Catalase, and HSP60–Apoferritin conditions. Because ROC curves plot the true positive rate against the false positive rate across all possible decision thresholds, they provide a threshold-independent view of a classifier’s underlying discriminatory capacity.

Finally, we report the abundance log ratio (ALR) of the minority species, which is defined as

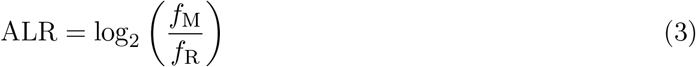

where *f*_M_ is the abundance predicted by the model for the minority species and *f*_R_ is the reference abundance of the minority species. This metric provides a quantitative measure of the difference between predicted and actual abundances.

### 2.5 Experimental Setup and Validation

To validate the effectiveness of our approach on imbalanced data, we designed a series of controlled experiments using three structurally distinct and well-characterized protein complexes: Catalase (tetrameric, 240 kDa)(Slowik *et al*., 2025), Apoferritin (octahedral, 480 kDa)(Kopylov *et al*., 2019), and HSP60 (heptameric double-ring, 800 kDa)(Clare *et al*., 2012). These proteins provide a range of symmetries and molecular weights, representing a tractable challenge for differential analysis.

#### 2.5.1 Sample Preparation and Mixing Ratios

For each protein, we prepared a stock solution with a mass concentration of 1 mg mL^*−*1^. HSP60 stock solution was prepared with a buffer of 50 mm Tris, 10 mm KCl, 10 mm MgCl_2_, pH 7.5. Apoferritin and Catalase stock solutions were prepared with a buffer of 50 mm Tris, 25 mm NaCl, pH 7.5. We prepared pairwise mixtures of each protein pair at a constant total protein concentration of 1 mg mL^*−*1^. These conditions provided the source particle data from which benchmark datasets at finer ratio increments were later derived by computational subsampling (see Data Processing).

For each mixing condition, we prepared two Quantifoil holey carbon grids (R 2/1 Cu 200 and 300 mesh) using a Vitrobot Mark IV (FEI/Thermo Fisher) with humidity 90 %, temperature 18 ^°^C, and blotting time 3.5 s. Grids were screened on a Glacios 200 kV transmission electron microscope (Thermo Fisher Scientific). The grid with more uniform ice thickness and superior particle distribution was selected from each pair for high-resolution data collection. However, since particle transfer was found to be markedly non-linear, we did not use the grids for quantitative analysis of the relative abundances of the species in the mixtures. Instead, we used computational subsampling to generate benchmark datasets with exact class ratios (see 2.5.3).

#### 2.5.2 Cryo-EM Data Acquisition

Data collection was performed on a Titan Krios G4 transmission electron microscope (Thermo Fisher Scientific) operating at 300 kV. The system was equipped with a Selectris X energy filter and a Falcon 4i direct electron detector. For each selected grid, we collected data at a nominal magnification of 165,000x, resulting in a calibrated pixel size of 0.73 Å. The total electron dose was set to 45 e^*−*^*/*Å^2^, fractionated over 64 frames to allow for motion correction. Defocus values were varied between *−*0.5 µm and *−*2.5 µm to ensure a continuous transfer of information across spatial frequencies. Automated data acquisition was managed via EPU software, targeting approximately 4000 micrographs per grid to ensure that minority species yielded enough particles to enable processing in cryoSPARC, the benchmark comparison.

#### 2.5.3 Data Processing and Analysis

Motion correction and dose weighting were performed using cryoSPARC Patch Motion Correction. CTF estimation was conducted with cryoSPARC PatchCTF. Particle picking was performed using a pre-trained Topaz model (ResNet 16 32 units) with estimated particle diameter 160 Å and extraction threshold 0, followed by extraction of particles with a box size of 360 px and downsampling to 128 px (2 px Å^*−*1^).

One round of cryoSPARC heterogeneous reconstruction was performed with 4 classes, followed by 2D classification per class. The results were curated manually to generate experimental reference labels. Particles from all conditions were pooled into a single labelled stack. This dataset was split into train/validation and test sets, with train/validation comprising 70 000 particles and the test set comprising 100 000 particles. The empty-pick class comprises 40 000 picks in the train/validation set and 50 000 picks in the test set. The train/validation set was used in one case (see Results 3.2), while the test set was reserved for evaluating the performance of CryoConvNeXt and of cryoSPARC for all conditions. The benchmark conditions were generated by subsampling pairs of species from the test set. The majority species and the empty class were kept at 100 000 and 50 000 samples respectively, while the minority species was subsampled from equal representation (1:1) to a 16-fold reduction (1:16).

To evaluate the performance of cryoSPARC (Punjani *et al*., 2017), heterogeneous reconstruction was performed for each benchmark condition. CryoSPARC receives the same 3D maps used as input to the TEM-simulator. From these, volumes are derived that serve as starting points for the particles present in the mixture (and an empty volume). CryoSPARC takes as input the experimental test set, or a subset thereof. The resulting class assignments were compared to the reference labels to calculate the *F*_1,macro_ score and ROC curves for each class. The Hydra algorithm (Levy *et al*., 2024*b*), which models compositional heterogeneity as a discrete mixture of neural fields, was evaluated on the HSP60–Catalase 16:1 condition (156 250 particles). Poses and CTF parameters were extracted from cryoSPARC heterogeneous reconstruction outputs. The model was configured with three classes, autodecoder-based conformation estimation, and trained for 50 SGD epochs (batch size 8) on a single 10 GB GPU over 24 h. CryoConvNeXt was evaluated by first training on the simulated dataset and then classifying the experimental test set, or a subset thereof, with performance metrics determined in the same way as with cryoSPARC for direct comparison. Self-training and training on experimental data were evaluated identically, apart from the changes to the training sets.

## 3 Results

All performance measurements reported below were obtained from experimentally collected cryo-EM particles. The classification results both from cryoSPARC heterogeneous reconstruction and from CryoConvNeXt are compared against manually curated reference labels. CryoConvNeXt was evaluated in three configurations differing only in training data: (i) simulated projections alone, (ii) simulation-based training followed by self-training from unlabelled experimental particles, and (iii) training on an independent, labelled experimental dataset. CryoSPARC was evaluated on the same experimental test particles. Benchmark conditions at different class ratios and particle numbers were generated by computational subsampling of the labelled test set: physical mixture ratios cannot be precisely controlled on holey carbon grids due to species-dependent particle transfer efficiency, and per-particle reference labels are required for precise performance evaluation.

We benchmarked all CryoConvNeXt configurations against cryoSPARC on binary mixtures and assessed per-class discrimination by ROC analysis. We then evaluated robustness to particle number and class imbalance, extended the benchmark from binary to ternary mixtures, and evaluated the abundance log-ratio (ALR) as a complementary metric for quantification accuracy.

### 3.1 Baseline Performance

For balanced datasets, the *F*_1,macro_ score is 0.75 for CryoConvNeXt trained on simulated data and 0.74 for cryoSPARC heterogeneous reconstruction, demonstrating that the model is competitive even in the scenario most favourable to cryoSPARC.

Per-class ROC curves for the balanced (1:1) Apoferritin–Catalase, HSP60–Catalase, and HSP60– Apoferritin pairs are provided in Appendix 5.3 (Figures 7, 8, and 9). Both CryoSPARC and CryoConvNeXt trained on simulated data achieve an AUC of 0.88 or higher for all protein classes, confirming that both methods can discriminate between species at balanced mixing ratios. CryoSPARC achieves higher AUC for the empty class, as expected: ice crystals and other contaminants present in experimental data are absent from the simulation and therefore not learnt by CryoConvNeXt.

### 3.2 Self-Learning and Training on Experimental Data

To partially bridge the simulation-to-real gap, we employed a self-training strategy (see Methods 2.3), which resulted in an improved *F*_1,macro_ score of 0.80 (vs. 0.75 before). Self-training requires no additional labelled data (it merely reuses its own classification output), is fully automated, and represents the highest simulation-to-real transfer learning performance achieved with CryoConvNeXt.

Simulation-to-real transfer learning is challenging and clearly a limiting factor in performance. To establish an upper bound on the performance of the CryoConvNeXt architecture under optimal conditions, we trained on a separate, independently curated experimental dataset and achieved an *F*_1,macro_ score of 0.91. This suggests that further improvements in simulation fidelity or in selftraining strategy can yield substantial performance boosts. The results are summarized in Table 1.

**Table 1:** Performance comparison of different training strategies for balanced classes. CryoConvNeXt trained on experimental data achieved the highest *F*_1,macro_ score. While CryoConvNeXt trained on simulated data performs on par with cryoSPARC heterogeneous reconstruction, self-training substantially outperforms cryoSPARC.

| Model | $F_{1,\text{macro}}$ |
| --- | --- |
| Trained on simulated data | 0.7549 |
| Trained on simulated data with self-training | 0.8010 |
| Trained on experimental data | 0.9129 |
| CryoSPARC heterogeneous reconstruction | 0.7407 |

### 3.3 Size and ratio of mixed-protein pairs

We subsampled the experimental reference test set to generate mixed protein pairs spanning a range of total particle counts and species ratios (Methods 2.5.3). Subsequently, we evaluated the performance of cryoSPARC heterogeneous reconstruction and CryoConvNeXt on these pairs, using the minority-species *F*_1_ score as the primary evaluation metric (Figure 2).

**Figure 1:**
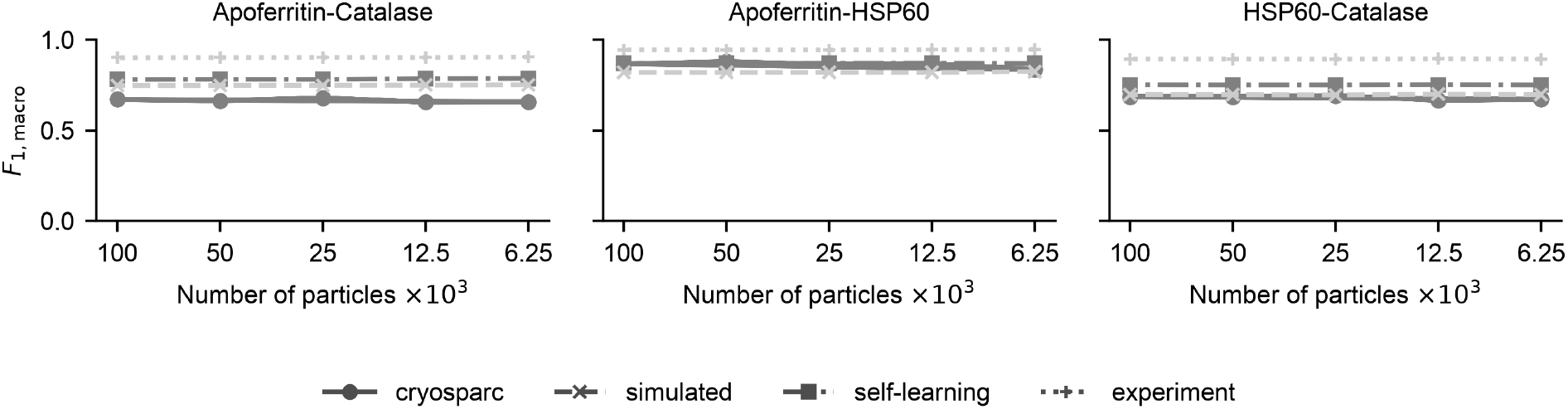
Balanced scenario: *F*_1,macro_ score for mixed-protein pairs is shown against the number of particles per class. The line style denotes the method: solid = cryoSPARC, dashed = CryoConvNeXt trained on simulated data, dash-dot = CryoConvNeXt trained with self-training, dotted = CryoConvNeXt trained on experimental data. Performance for all methods remains stable as the number of particles decreases.

**Figure 2:**
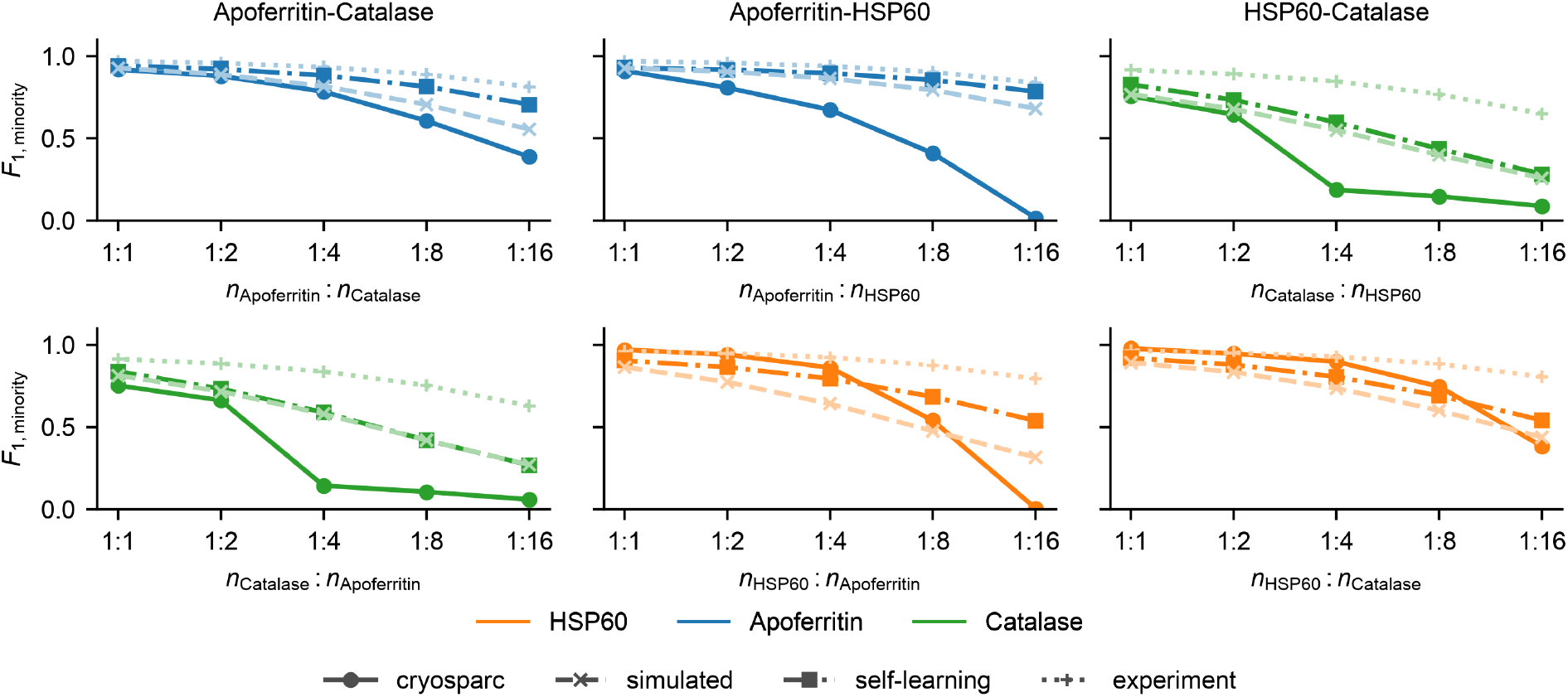
Minority-species *F*_1_ score as a function of species ratio for all pairwise protein mixtures. Each column corresponds to a protein pair (Apoferritin–Catalase, Apoferritin–HSP60, HSP60– Catalase) and each row corresponds to a minority species. Line style indicates method: solid = cryoSPARC heterogeneous reconstruction, dashed = CryoConvNeXt trained on simulated data, dash-dot = CryoConvNeXt with self-training, dotted = CryoConvNeXt trained on experimental data. CryoSPARC’s performance degrades sharply as the minority fraction decreases, whereas CryoConvNeXt’s performance decreases far more slowly.

While cryoSPARC’s classification performance remains robust against a uniform reduction in data volume under balanced conditions, we observed that it deteriorated severely as class imbalance increased (Figure 2). For instance, in the Apoferritin–Catalase pair at a 1:1 ratio (100 000 particles), cryoSPARC achieves an *F*_1_ score of approximately 0.65 for Catalase; however, when Catalase becomes the minority species at 6250 particles (ratio 1:16), its *F*_1_ score drops close to zero. This highlights a fundamental limitation of conventional multi-reference 3D classification workflows when extracting rare structural populations. Because these iterative algorithms rely on global volume updates driven by data density, the alignment signal of the minority class is easily marginalized by the dominant species.

In contrast, CryoConvNeXt demonstrates superior resilience to class imbalance. Because Cryo-ConvNeXt evaluates each particle image independently, its feature-extraction capabilities are fundamentally decoupled from the global dataset composition.

Nevertheless, CryoConvNeXt does exhibit a gradual, less severe decline in minority-species *F*_1_ scores as the minority fraction decreases. Because CryoConvNeXt maintains a stable, independent error rate per image, evaluating it on a dataset dominated by a majority species causes a fixed percentage of majority-class false positives to progressively outnumber the shrinking pool of true positives. This disproportionately penalizes precision, resulting in the observed drop in the minority *F*_1_ score despite the underlying model stability.

The recall curves (Figure 3) reveal a qualitative difference between the two methods. In several conditions, cryoSPARC undergoes a catastrophic collapse in minority-species recall as the majorityto-minority ratio increases, effectively becoming blind to the minority class entirely. CryoConv-NeXt’s recall, by contrast, remains stable across all tested ratios, and only precision decreases, for the reasons discussed above. This distinction has practical consequences. A stable recall means that minority species are reliably detected even at low abundance; a core requirement for quantitative analysis. Precision losses at high ratios are, in principle, addressable by improving the simulation fidelity, by expanding the training set, or by post hoc processing (i.e. using CryoConvNeXt as a step in a larger processing pipeline). However, complete collapse of recall, as occurs with cryoSPARC, cannot be recovered by any post hoc adjustment.

**Figure 3:**
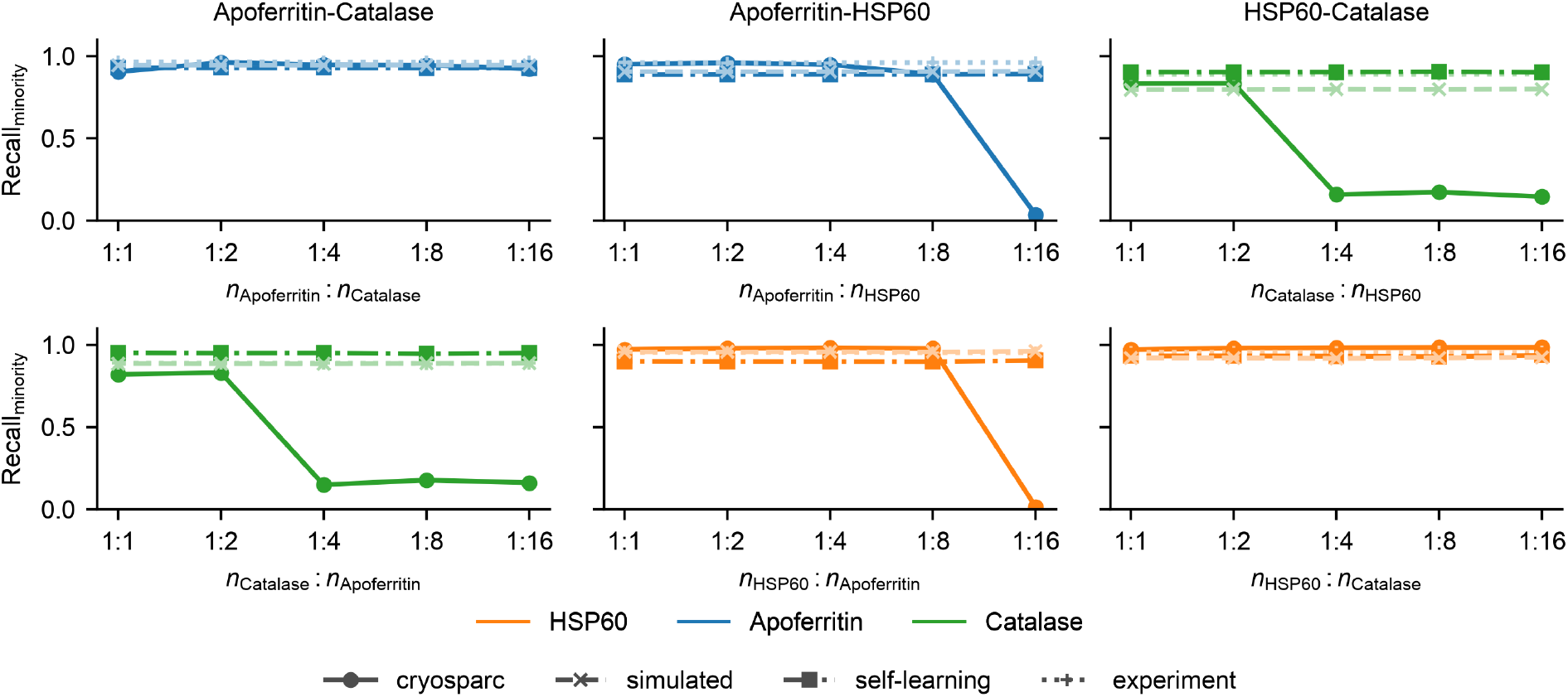
Minority-species recall for mixed-protein pairs. Each column corresponds to a protein pair (Apoferritin–Catalase, Apoferritin–HSP60, HSP60–Catalase) and each row corresponds to a minority species. The line style denotes the method: solid = cryoSPARC, dashed = CryoConvNeXt trained on simulated data, dash-dot = CryoConvNeXt trained with self-training, dotted = CryoConvNeXt trained on experimental data. Catalase suffers recall collapse at a ratio of 1:4; Apoferritin and HSP60 also collapse when paired together. CryoConvNeXt recall remains stable in all conditions.

**Figure 4:**
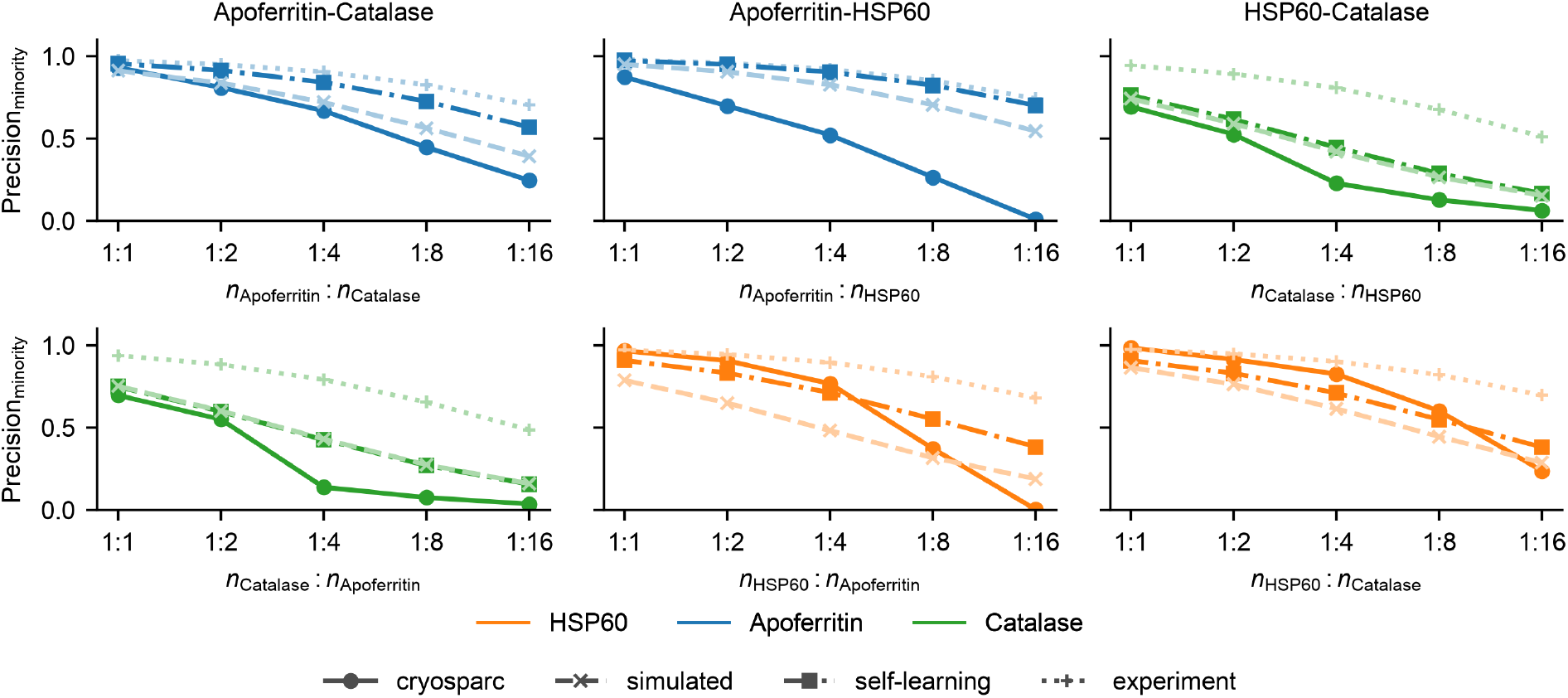
Minority-species precision for mixed-protein pairs. Each column corresponds to a protein pair (Apoferritin–Catalase, Apoferritin–HSP60, HSP60–Catalase) and each row corresponds to a minority species. The line style denotes the method: solid = cryoSPARC, dashed = CryoConv-NeXt trained on simulated data, dash-dot = CryoConvNeXt trained with self-training, dotted = CryoConvNeXt trained on experimental data. All classifications show decreased precision with increasing imbalance, to different degrees.

The ALR metric (Figure 5) provides a complementary perspective on quantification accuracy. For most conditions, CryoConvNeXt achieves smaller |ALR| than cryoSPARC, indicating more accurate abundance estimates. An exception arises when Catalase is the minority species: here cryoSPARC can appear to match or outperform CryoConvNeXt on ALR. However, Figure 3 shows that cryoSPARC undergoes catastrophic recall collapse in exactly these conditions, and precision is similarly low. The apparent accuracy on ALR is coincidental: when *f*_*R*_ is small, a near-zero *f*_*M*_ from recall collapse combined with a fortuitous number of false positives can yield a small |ALR| — not because abundance is correctly estimated, but because the errors happen to mostly cancel each other out for low abundance values. This illustrates why ALR must be interpreted alongside recall and precision: a low |ALR| is only meaningful when the species is detected and the detections are accurate.

**Figure 5:**
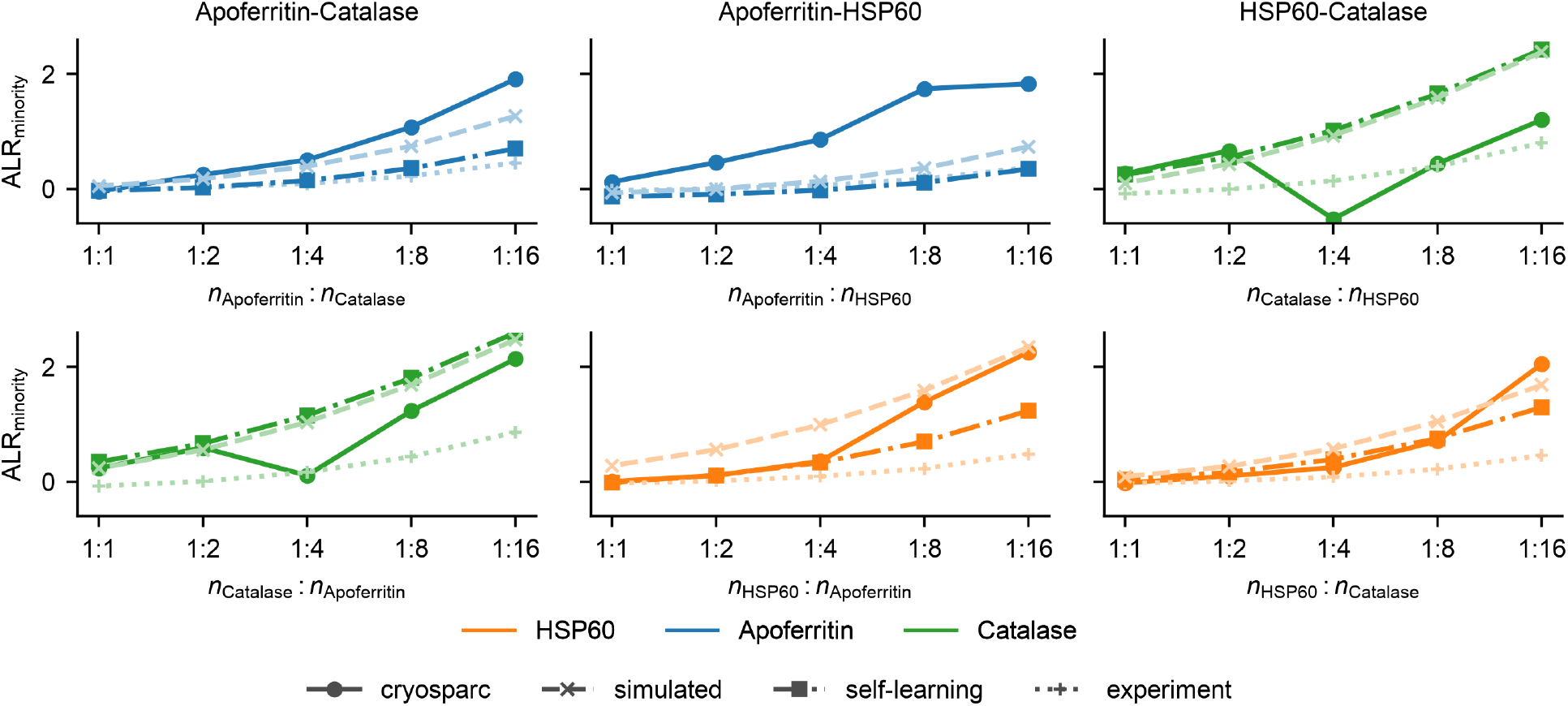
Minority-species ALR_minority_ as a function of species ratio for all pairwise protein mixtures. Each column corresponds to a protein pair (Apoferritin–Catalase, Apoferritin–HSP60, HSP60– Catalase) and each row corresponds to a minority species. Line style indicates method: solid = cryoSPARC heterogeneous reconstruction, dashed = CryoConvNeXt trained on simulated data, dash-dot = CryoConvNeXt with self-training, dotted = CryoConvNeXt trained on experimental data.

**Figure 6:**
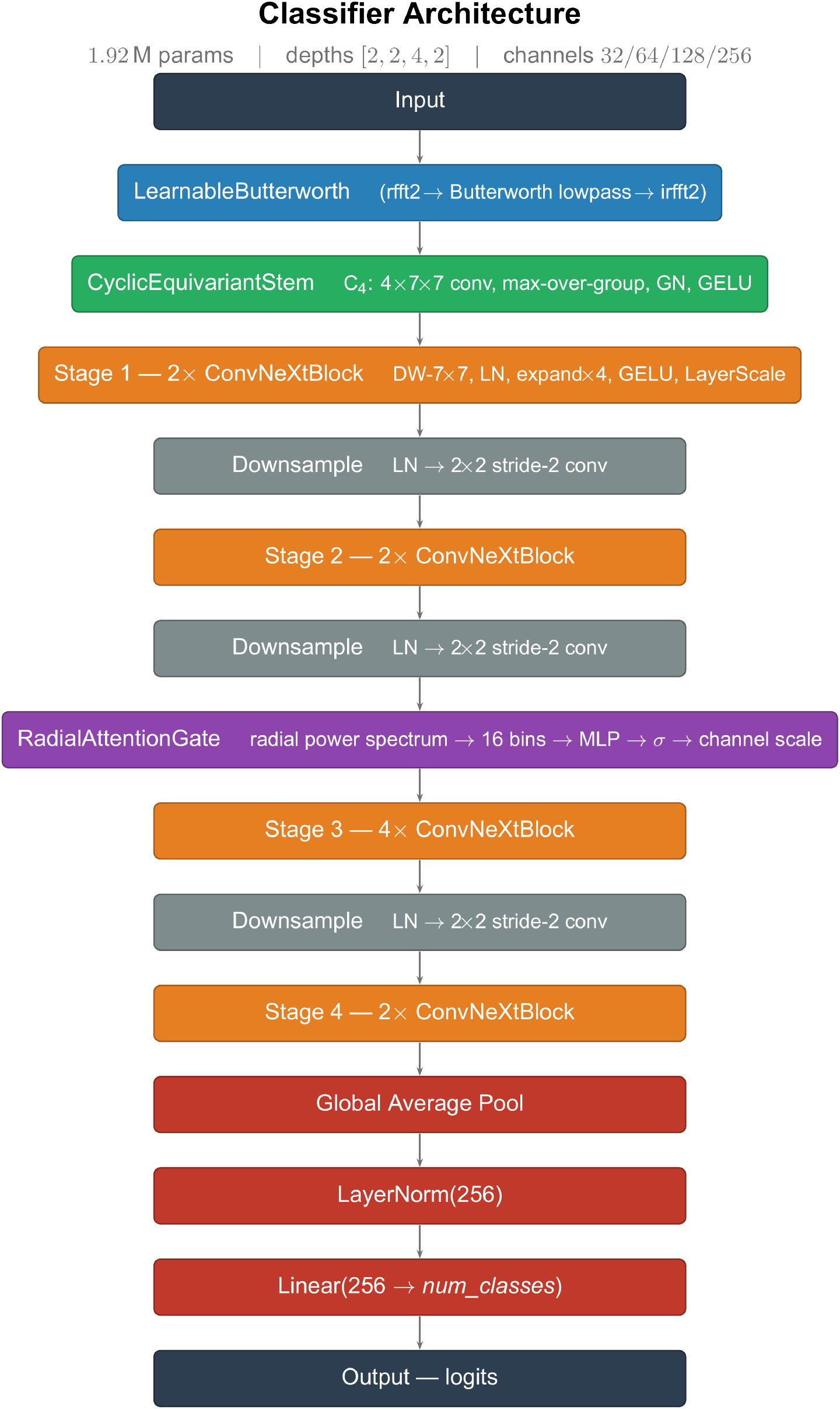
CryoConvNeXt architecture.

**Figure 7:**
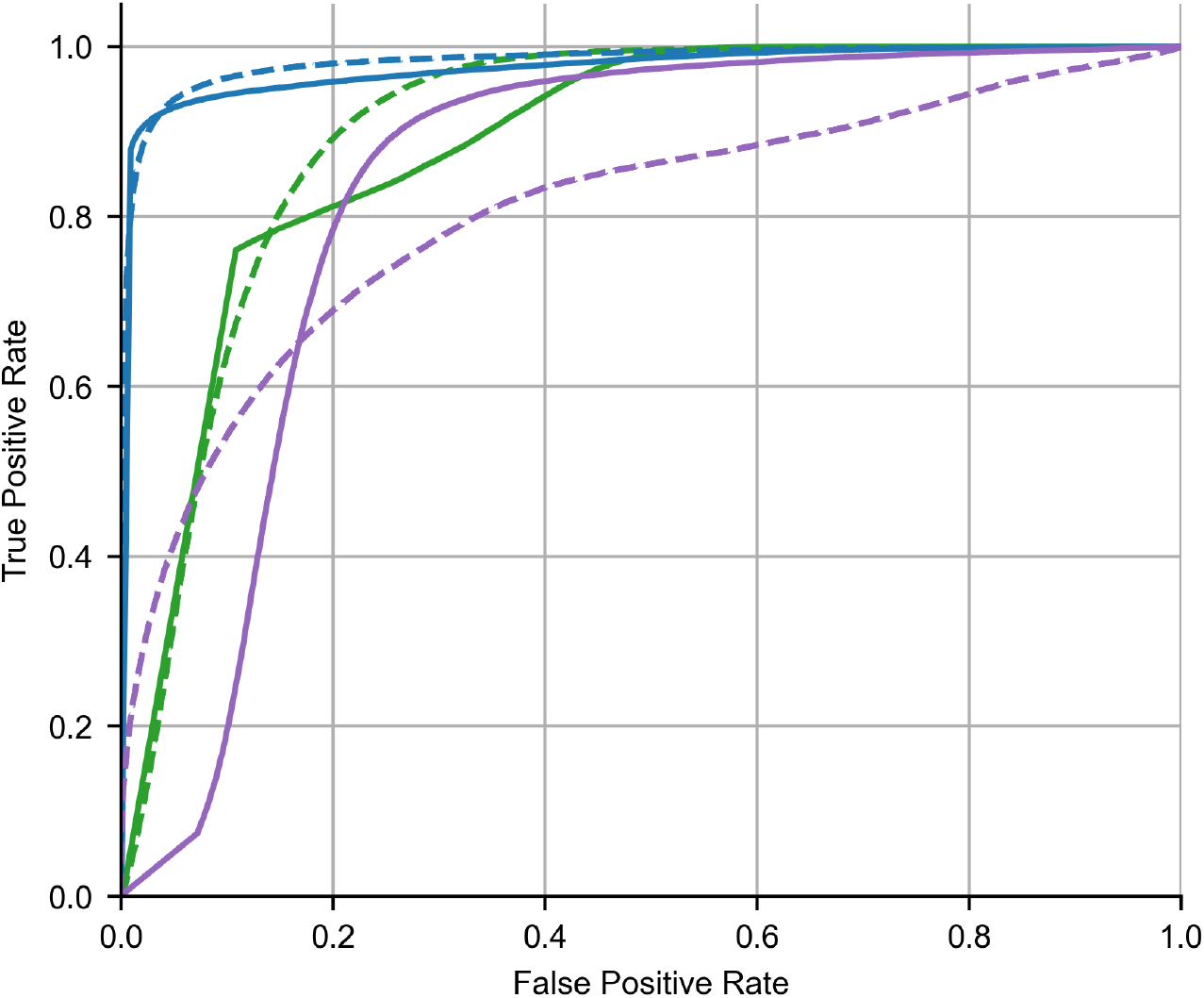
Multiclass ROC (one-vs-rest) curves for the Apoferritin–Catalase pair. The line style denotes the method: solid = cryoSPARC, dashed = CryoConvNeXt trained on simulated data.

**Figure 8:**
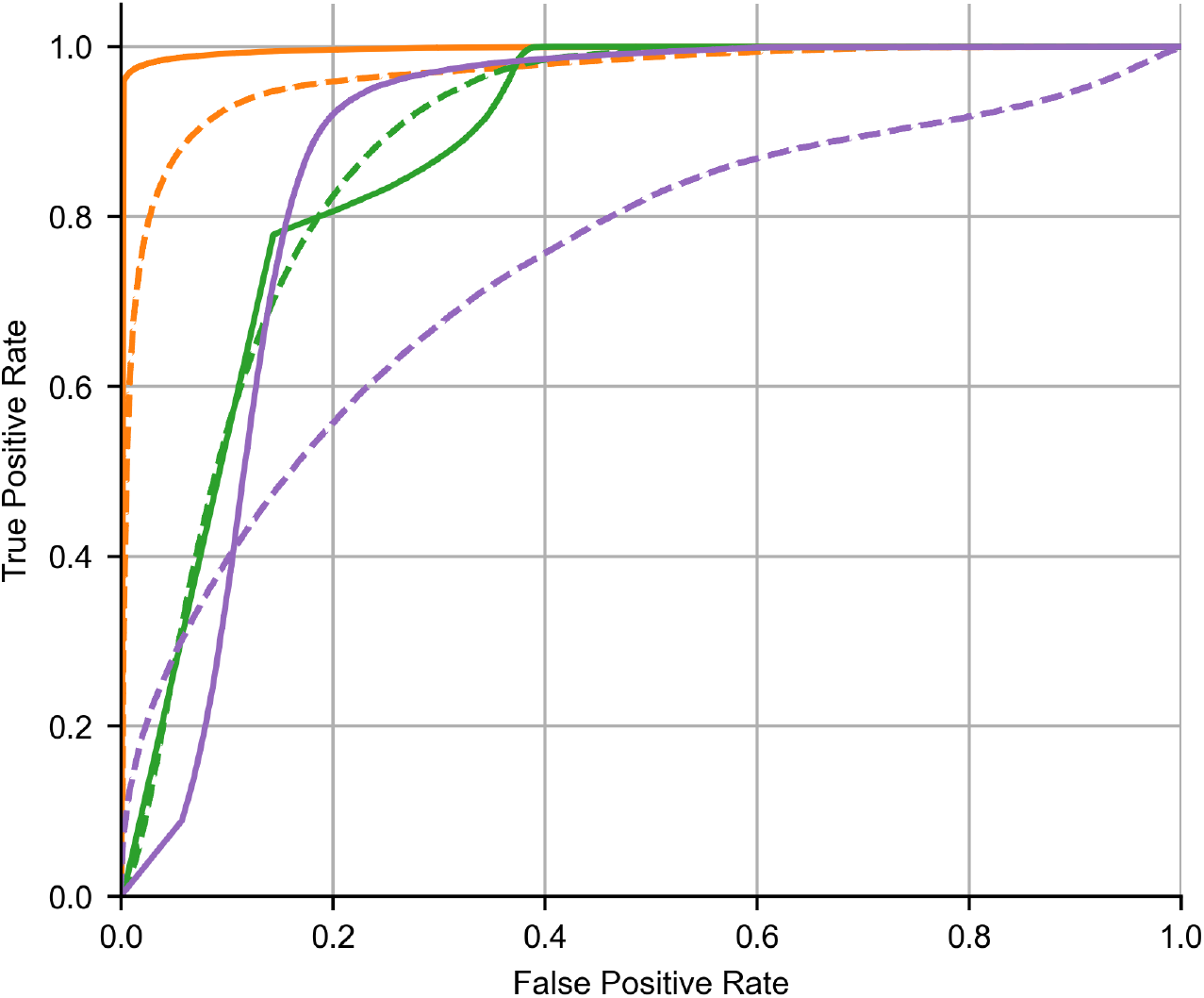
Multiclass ROC (one-vs-rest) curves for the HSP60–Catalase pair. The line style denotes the method: solid = cryoSPARC, dashed = CryoConvNeXt trained on simulated data.

**Figure 9:**
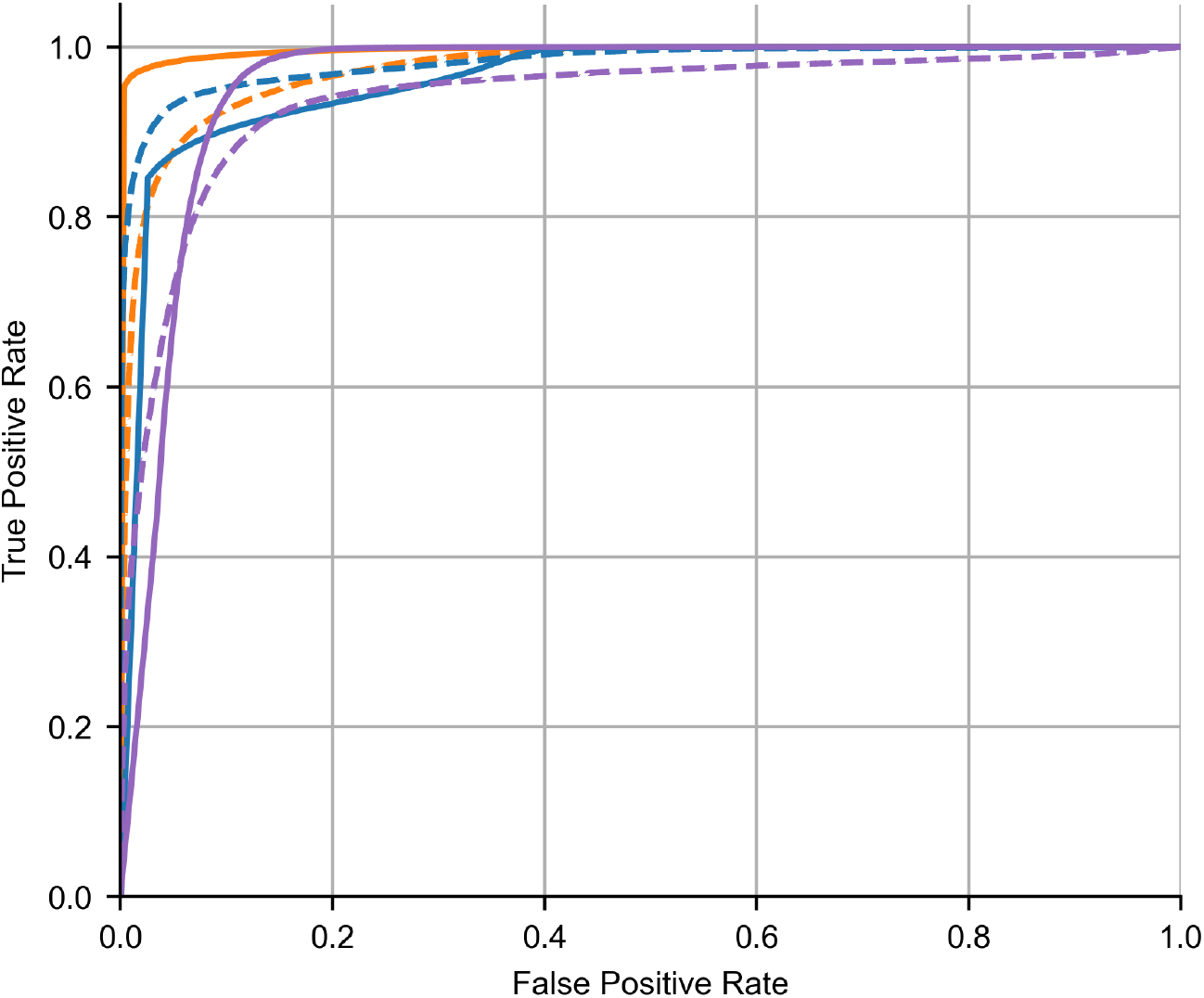
Multiclass ROC (one-vs-rest) curves for the HSP60–Apoferritin pair. The line style denotes the method: solid = cryoSPARC, dashed = CryoConvNeXt trained on simulated data.

The Hydra algorithm (Levy *et al*., 2024*b*) achieved an *F*_1,macro_ of 0.30 on the HSP60–Catalase 16:1 condition, with a minority-species (Catalase) *F*_1_ of 0.10 and ALR = 3.01, indicating that the model assigns approximately 8*×* more particles to the Catalase class than are present.

### 3.4 Number of Proteins

We extended the benchmark to ternary mixtures. The results are summarized in Table 2. CryoConvNeXt trained on simulated data achieved an *F*_1,macro_ score of 0.75, the same as for binary mixtures. The model trained on simulated data with self-training achieved an *F*_1,macro_ score of 0.78, compared to 0.80 for binary mixtures. CryoSPARC heterogeneous reconstruction achieved an *F*_1,macro_ score of 0.74, the same as for binary mixtures. The performance is comparable to the binary mixtures, indicating that the approach scales well with increasing sample complexity.

**Table 2:** Performance comparison of different training strategies on a ternary mixture. All training strategies achieve comparable performance to binary mixtures.

| Model | $F_{1,\text{macro}}$ |
| --- | --- |
| Trained on simulated data | 0.7477 |
| Trained on simulated data with self-training | 0.7838 |
| Trained on experimental data | 0.9049 |
| CryoSPARC heterogeneous reconstruction | 0.7358 |

**Table 3:** Simulation parameters used for the generation of synthetic cryo-EM images.

| Parameter | Value |
| --- | --- |
| Microscope Voltage | 300 kV |
| Energy Spread | 0.1 V |
| Dose | $4500 \text{ e}^-/\text{nm}^2$ |
| Physical Pixel Size | $14 \mu\text{m}$ |
| Gain | 1 |
| DQE | 0.9 |
| $MTF_a$ | 0.5 |
| $MTF_b$ | 0.5 |
| $MTF_c$ | 0 |
| $MTF_\alpha$ | 7 |
| $MTF_\beta$ | 20 |
| $MTF_p$ | 4 |
| $MTF_q$ | 2 |
| $C_s$ | 2.7 mm |
| $C_c$ | 2.0 mm |
| Aperture Size | $100 \mu\text{m}$ |
| Focal Length | 3.4 mm |
| Condensor Aperture Angle | 0.005 mrad |
| Defocus Nonsystemic Error | $0.02 \mu\text{m}$ |
| Magnification | 191,000x |
| Defocus Range | $-0.5 \mu\text{m}$ to $-2.5 \mu\text{m}$ |

## 4 Discussion

We demonstrate that a deep-learning classifier trained on simulated cryo-EM projections can reliably recover minority species from experimentally collected data, validated against manually curated perparticle reference labels. On balanced binary mixtures, CryoConvNeXt with self-training achieves an *F*_1,macro_ score of 0.80, outperforming cryoSPARC heterogeneous reconstruction evaluated under identical conditions (0.74). This advantage increases rapidly with class imbalance, the scenario most relevant to endogenous biological samples. CryoSPARC undergoes catastrophic minority-class recall collapse in four of six binary comparisons, becoming effectively blind to the minority species, while CryoConvNeXt’s recall remains stable across all tested ratios.

The performance difference has a straightforward explanation rooted in the different objectives of the two methods. CryoSPARC heterogeneous reconstruction was designed to optimise a reconstruction objective; assigning particles to classes is a secondary output, and it has no mechanism for upweighting minority species. Hydra (Levy *et al*., 2024*b*), which models compositional heterogeneity as a mixture of neural fields, similarly failed to recover the minority species at 1:16, consistent with the view that the limitation is shared by reconstruction-based objectives under class imbalance. CryoConvNeXt, optimised for a classification objective with label-smoothing cross-entropy loss, is better suited to this task by design.

The gap between CryoConvNeXt without self-training (*F*_1,macro_ = 0.75) and the same architecture trained on experimental data (*F*_1,macro_ = 0.91) reflects the challenges of simulation-to-real transfer learning: while the simulation models Poisson and detector noise calibrated to the Falcon 4i MTF, it cannot reproduce sample-specific factors such as ice contamination, salt-induced contrast variations, unfolded proteins, and grid-to-grid variability. The model trained on experimental data reveals an upper bound on the performance achievable with this architecture. Self-training partially bridges the simulation-to-real gap by incorporating high-confidence experimental predictions as pseudolabels and improving performance to *F*_1,macro_ = 0.80, without consulting any reference labels. The approach is transductive: the model adapts to the target data distribution before inference. This matches the intended use case where self-training is run on the dataset being classified. However, self-training preferentially selects particles already well-predicted by the simulation, leaving out-of-distribution particles underrepresented. Future work could target this residual gap directly through active learning strategies that specifically seek out such hard cases.

Several limitations of the current benchmark should be noted. The three proteins tested, Catalase (240 kDa, tetrameric), Apoferritin (480 kDa, octahedral), and HSP60 (800 kDa, heptameric double-ring), differ substantially in size and symmetry, making them a relatively favourable classification task. Performance on mixtures of similarly sized proteins, proteins with less distinct projection shapes, or mixtures of more than three components, remains to be evaluated. The approach requires a reference 3D density map for each species and these maps must carry physically meaningful values, as they are the basis for simulating projection images rather than refinement initialisations.

This constrains applicability to species for which structures already exist or can be predicted with confidence. The same limitation applies across supervised classification approaches generally, and becomes less restrictive as structural databases continue to expand. CryoConvNeXt scales to three-protein mixtures with minimal performance loss (simulation-trained: *F*_1,macro_ = 0.75; self-training: 0.78; experimentally trained: 0.90), though the slight reduction in self-training performance relative to binary mixtures (0.80) is statistically significant (95 % bootstrap CI: [0.783, 0.785] vs. [0.818, 0.820]). With three classes, confidence is distributed more broadly, increasing the probability that a high-confidence prediction is nonetheless incorrect; erroneous pseudo-labels then partially counteract the benefit of incorporating experimental data.

Quantitative visual proteomics of endogenous samples requires three capabilities in concert: low-loss microfluidic sample preparation, a grid support providing linear and species-independent particle transfer, and accurate classification of the resulting particle images. The first has been demonstrated using microfluidic vitrification workflows (Arnold *et al*., 2017; Kemmerling *et al*., 2013; Arnold *et al*., 2018). Linear particle transfer on continuous carbon support films has been established for negative-stain preparations (Giss *et al*., 2014); verifying this under cryo-EM conditions and adapting the simulator to model the continuous carbon background remain open tasks, while the classification framework itself is grid-independent. The present work addresses the third requirement, enabling the recovery of minority species at low abundance that conventional heterogeneous reconstruction overlooks. With all three capabilities in place, two applications become particularly compelling: characterising the full complement of macromolecular species across biological conditions, and quantifying the stoichiometry of transient hetero-complexes, where composition encodes functional state and cannot be inferred from structure alone.

## Acknowledgements

We thank Rémi Ruedas (BioEM Lab, University of Basel) for his help with the sample handling. We thank Colin W. Glass for helpful discussions.

This work was supported by the Swiss National Foundation (SNF project 200020_192190).

Molecular graphics and analyses performed with UCSF ChimeraX, developed by the Resource for Biocomputing, Visualization, and Informatics at the University of California, San Francisco, with support from National Institutes of Health R01-GM129325 and the Office of Cyber Infrastructure and Computational Biology, National Institute of Allergy and Infectious Diseases.

## Data and Code Availability

The cryo-EM data generated in this study is currently being submitted to EMPIAR. The code for CryoConvNeXt will be made publicly available upon journal acceptance.

## Appendix

### 5.1 Simulation Details

The simulation is calibrated to match the experimental conditions, including the modulation transfer function (MTF) of the detector and the contrast transfer function (CTF) of the microscope.

#### 5.2 Architecture Diagram

#### 5.3 ROC Curves

